# Structural insight into TRPV5 channel function and modulation

**DOI:** 10.1101/434902

**Authors:** Shangyu Dang, Mark K. van Goor, YongQiang Wang, David Julius, Yifan Cheng, Jenny van der Wijst

**Affiliations:** Department of Biochemistry and Biophysics, University of California San Francisco, San Francisco, CA, USA; Department of Physiology, Radboud Institute for Molecular Life Sciences, Radboud university medical center, Nijmegen, The Netherlands; Department of Physiology, University of California, San Francisco, California, 94143, USA; Howard Hughes Medical Institute, University of California San Francisco, San Francisco, CA, USA

**Author notes:** Jenny van der Wijst, PhD Department of Physiology Radboud Institute for Molecular Life Sciences Radboud university medical center P.O. Box 9101, 6500 HB, Nijmegen, The Netherlands; Yifan Cheng, PhD Department of Biochemistry & Biophysics University of California San Francisco Mission Bay, Genentech Hall, MC-2240 600 16th Street, Room S472D San Francisco, CA 94158-2517, USA; David Julius, PhD Department of Physiology University of California San Francisco Mission Bay, Genentech Hall, MC-2140 600 16th Street, Room N272E San Francisco, CA 94158-2517, USA.

## Abstract

TRPV5 (transient receptor potential vanilloid) is a unique calcium-selective TRP channel that is essential for calcium homeostasis. TRPV5 and its close homologue TRPV6 do not exhibit thermosensitivity or ligand-dependent activation, unlike other TRPV channels, but are constitutively opened at physiological membrane potentials. Here, we report high resolution electron cryo-microscopy (cryo-EM) structures of truncated and full length TRPV5 in lipid nanodisc, as well as a TRPV5 W583A mutant structure and a complex structure of TRPV5 with calmodulin (CaM). These structures highlight and explain functional differences between the thermosensitive and calcium-selective TRPV channels. An extended S1-S2 linker folds on top of the channel that might shield it from modulation by extracellular factors. Resident lipid densities in the homologous vanilloid pocket are different from those previously found in TRPV1, supporting a comparatively more rigid architecture of TRPV5. A ring of tryptophan residues (W583) at the bottom of the pore coordinates a density and mutation of W583 resultes in opening of the lower gate. Moreover, we provide structural insight into the calcium-dependent channel inactivation and propose a flexible stoichiometry for TRPV5 and CaM binding.

## Main

The transient potential receptor (TRP) family is one of the largest classes of ion channels, with diverse functionality. This is exemplified by the existence of channel-specific ligand activators, differences in ion selectivity, and mode of activation and permeation. TRPV5 is unique amongst all TRP channels because of its high selectivity for calcium over monovalent ions (P_Ca_/P_Na_ > 100:1) (1). Together with its close homolog TRPV6, they are also different from other members of the vanilloid subfamily (TRPV1-4) and are not considered to be thermosensitive or ligand-activated (2, 3). Physiologically, TRPV5 and TRPV6 serve as apical entryways into epithelial cells that line parts of the gut and nephron, initiating transcellular calcium transport pathways that help fine-tune serum calcium levels (4). Disturbances in the body’s calcium balance can cause major health problems, including neurological and cardiac aberrations, stone formation, and bone disorders.

In addition to the specific calcium-selectivity, the functional hallmark of TRPV5 is a pronounced inward rectification of the current-voltage relationship, compared to an outward rectification for most other TRP channels. This allows TRPV5 to move calcium ions into the cell at negative membrane potential, resulting in significant calcium permeation at physiological membrane potentials. Furthermore, TRPV5 channels exhibit calcium-dependent feedback regulation that includes fast inactivation and slow current decay (5). This inhibition is likely controlled by the concentration of calcium locally near the cytoplasmic side of the channel, and prevents excessive calcium influx. Part of the calcium-mediated negative feedback mechanism functions via binding of calmodulin (CaM), a ubiquitous calcium-binding protein (6, 7).

To understand how the structure of the pore conveys extraordinary calcium selectivity and calcium-dependent channel inactivation, we set out to determine the structure of the epithelial calcium channel TRPV5. Here, we report structures of truncated and full length rabbit TRPV5 channels at nominal resolutions of 2.9 Å and 3.0 Å, respectively. In addition, we determined the structure of the TRPV5-CaM complex in LMNG detergent at a nominal resolution of 3.0 Å, as well as the TRPV5 W583A mutant as an open channel structure at 2.85 Å.

## Results

### Structure of TRPV5

We expressed and purified rabbit TRPV5 with a C-terminal truncation (1-660) as well as in full length channel (1-730). Proteins purified from both constructs were stable and monodisperse (Supplemental Fig. 1a-b). The truncated TRPV5 retains most of its functional domains, but lacks one of the postulated calmodulin-binding sites (701-730) that was shown to regulate TRPV5 channel activity in a calcium-dependent manner(6, 7). The truncated channel shows significant, but slightly less robust TRPV5-mediated calcium uptake compared to the full length protein (Supplemental Fig. 1c). We solubilized and purified both full length and truncated TRPV5 in detergent, followed by reconstitution into lipid nanodiscs for single particle cryo-EM studies. Electron micrographs of frozen hydrated nanodisc-reconstituted proteins show monodispersed particles without obvious preferred orientations (Supplemental Fig. 2a,b). Two-dimensional (2D) class averages calculated from selected particles of both samples showed clear features of both transmembrane and intracellular domains (Supplemental Fig. 2c), as well as a density corresponding to the nanodisc that surrounds the transmembrane core of the channel (Fig. 1a,b). To elucidate the structure and the gating mechanism of TRPV5, we determined a total of four 3D reconstructions, including truncated and full length TRPV5 in lipid nanodisc at overall resolutions of 2.9 Å and 3.0 Å, respectively (Fig. 1, Supplemental Fig. 2), as well as TRPV5-CaM complex in lauryl maltose neopentyl glycol (LMNG) detergent and full length protein with a W583A point mutation in lipid nanodisc to resolutions of 3.0Å and 2.85Å respectively (Supplemental Fig. 3,4). The resolutions and the qualities of all maps were sufficient to trace protein backbone and model side chains of most residues unambiguously (residues 27 to 638) (Fig. 1).

**Figure 1.**
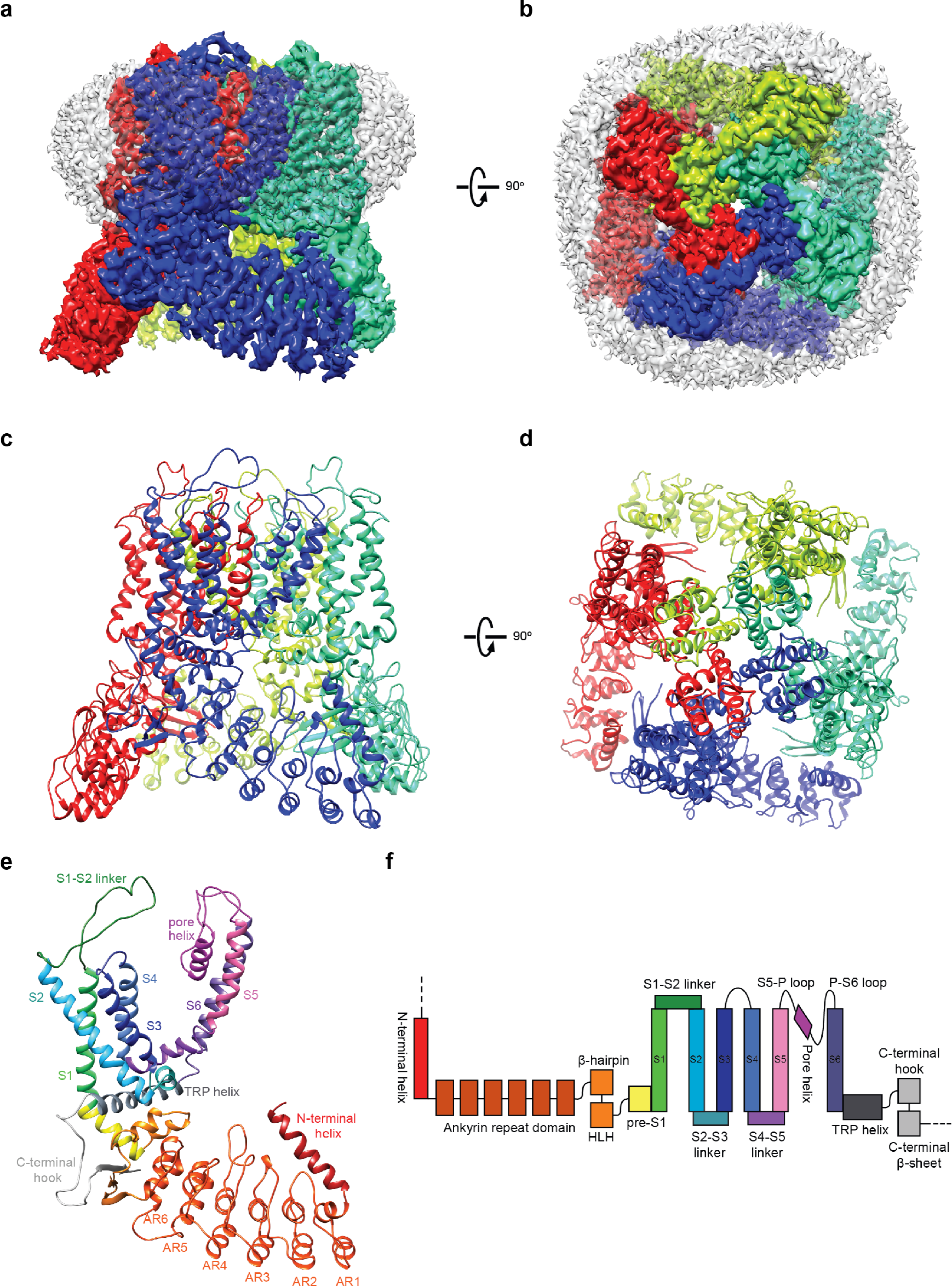
Cryo-EM structure of TRPV5. **a,b,** Side and top view of nanodisc-reconstituted TRPV5 660X at 2.9 Å resolution, with the subunits colored individually. The nanodisc density surrounds the transmembrane region of the channel (colored in white). **c,d**, Side and top view of the TRPV5 tetrameric complex, using the same subunit colors as panels **a,b**. **e,**Side view of a TRPV5 monomer with the functional domains colored individually. Structural elements are depicted as either ribbons, for alpha-helices, arrows, for beta-sheets and ropes for unordered loops. **f,**Domain organization of TRPV5, with domains colored as in panel e. The dashed line denotes a region that was not resolved in the structure but is present in the full-length channel.

In all of our TRPV5 structures in lipid nanodisc, many densities that can be attributed to annular lipids are well resolved, occupying crevasses between two neighboring units (Fig. 2a, Supplemental Fig. 6a,b). In addition, one lipid density is found within the lower segment of the S1-S4 domain. The location and shape of this lipid density is very similar to the one found in TRPV1, where it was interpreted as a phosphatidylcholine (8). Another lipid is found in a pocket akin to the vanilloid binding site in TRPV1, which is occupied by a phosphatidylinositol lipid in apo TRPV1; binding of a vanilloid compound in this pocket replaces the phosphatidylinositol and activates TRPV1 (8). In TRPV5, however, the shape of this lipid density is clearly not consistent with a phosphatidylinositol, but more likely represents an annular lipid (Fig. 2b). While we do not know the function of these two lipids bound to TRPV5, they form tight interactions with the protein and are likely endogenous lipids that remain tightly bound to the channel during the purification process.

**Figure 2.**
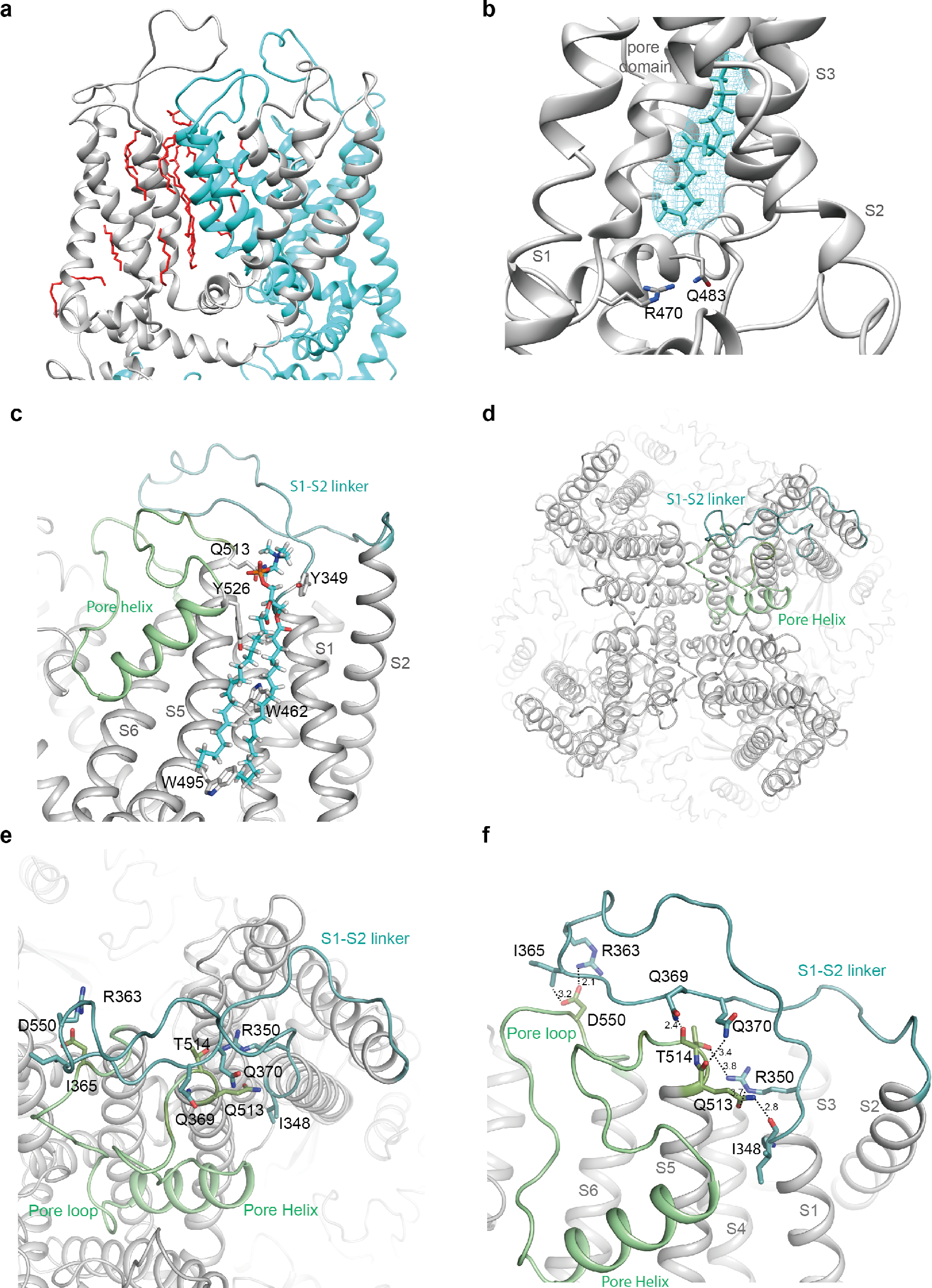
S1-S2 linker position in TRPV5. **a,** Expanded view of the lipid densities observed within the pore domain. Two monomers are shown and individually colored. Lipids are red. **b,** cZoom of the resident lipid in the vanilloid pocket. The density of the lipid is visualized with cyan mesh and the fitted acyl chain is colored cyan. The side chains of residues important in phosphatidylinositol-binding in TRPV1 are shown. **c,**Zoom of the interaction formed by the S1-S2 linker, annular lipid, and pore domain. The side chains of residues interaction with the lipid are shown. **d,**Top view of TRPV5 tetramer showing the positions of the S1-S2 linker (light teal) and S5-P-S6 domain (pale green). **e,f,** Zoom of the inter-subunit interface formed by the S1-S2 linker and the pore helix. Putative hydrogen bonds and electrostatic interactions are displayed as dashed lines. Side chains of interacting residues are shown as sticks for both and interatomic distances (**f**) between side chains are depicted.

Overall, TRPV5 closely resembles the structures of other TRPV subfamily members with its six transmembrane helices arranged in a domain-swapped architecture and its N-terminus dominated by an ankyrin repeat domain (ARD) (Fig. 1c-f) (9–14). Similar to TRPV6 (but not observed in TRPV1-4) is a helix preceding the first ankyrin repeat that forms a major interface between neighboring subunits at each corner of the channel. Moreover, an extended extracellular linker between S1 and S2 is present in TRPV5 (and TRPV6), but lacking in other TRPV members (Supplemental Fig. 6c,d, Supplemental Fig. 10). This linker interacts with the pore region of the neighboring subunit and contains the N-glycosylation site(s) N358 and N357 (Fig. 2d, Supplemental Fig. 6c,d) (15). It is stabilized by forming rather tight interactions through hydrogen bonds between the carboxyl group of D550 in the pore loop and R363 and I365 in the S1-S2 loop (Fig. 2e-f), a web of other hydrogen bonds and salt bridges between I348, R350, Q369, Q370 in S1-S2 loop and Q513, T514 in the S5-Pore loop, and some main chain interactions. Q513 is located at the top of the pore domain and seems to play a major role in interacting with the S1-S2 loop, by contributing to a number of hydrogen bonds and salt bridges with the S1-S2 loop (Fig. 2e-f). As one of the few TRPV5/TRPV6 specific regions, such interaction may constrain the movement of the selectivity filter. Another interesting structural feature for TRPV5 and 6 is a C-terminal helix, uniquely observed in our CaM-bound structure, that will be discussed in more detail below.

### Ion permeation pathway

Similar to other TRPV ion channels, the ion permeation pathway of TPRV5 has two main constriction points, or gates. The upper gate is formed by D542, which is located in the lower part but not the bottom of the pore loop. Above the upper gate is a large cavity formed by the upper part of the pore loop, which is stabilized by the S1-S2 linker loop. This conformation is somewhat different from either the funnel shaped configuration of the outer pore in the apo structure of TRPV1, where the upper gate is in a closed conformation, or the parallel conformation observed in an open conformation when in complex with the agonist spider toxin, DkTx(16). In our structure, there is a density located in the middle of the pore in between the side chains of D542 (Supplemental Fig. 7a), where similar densities are found in the same location in both half maps. The distance between the side chains of opposite D542 residues is about 5Å (Fig. 3a), sufficient to coordinate a cation. Thus, this density likely corresponds to a coordinated cation, consistent with D542 serving as a cation selectivity filter.

**Figure 3.**
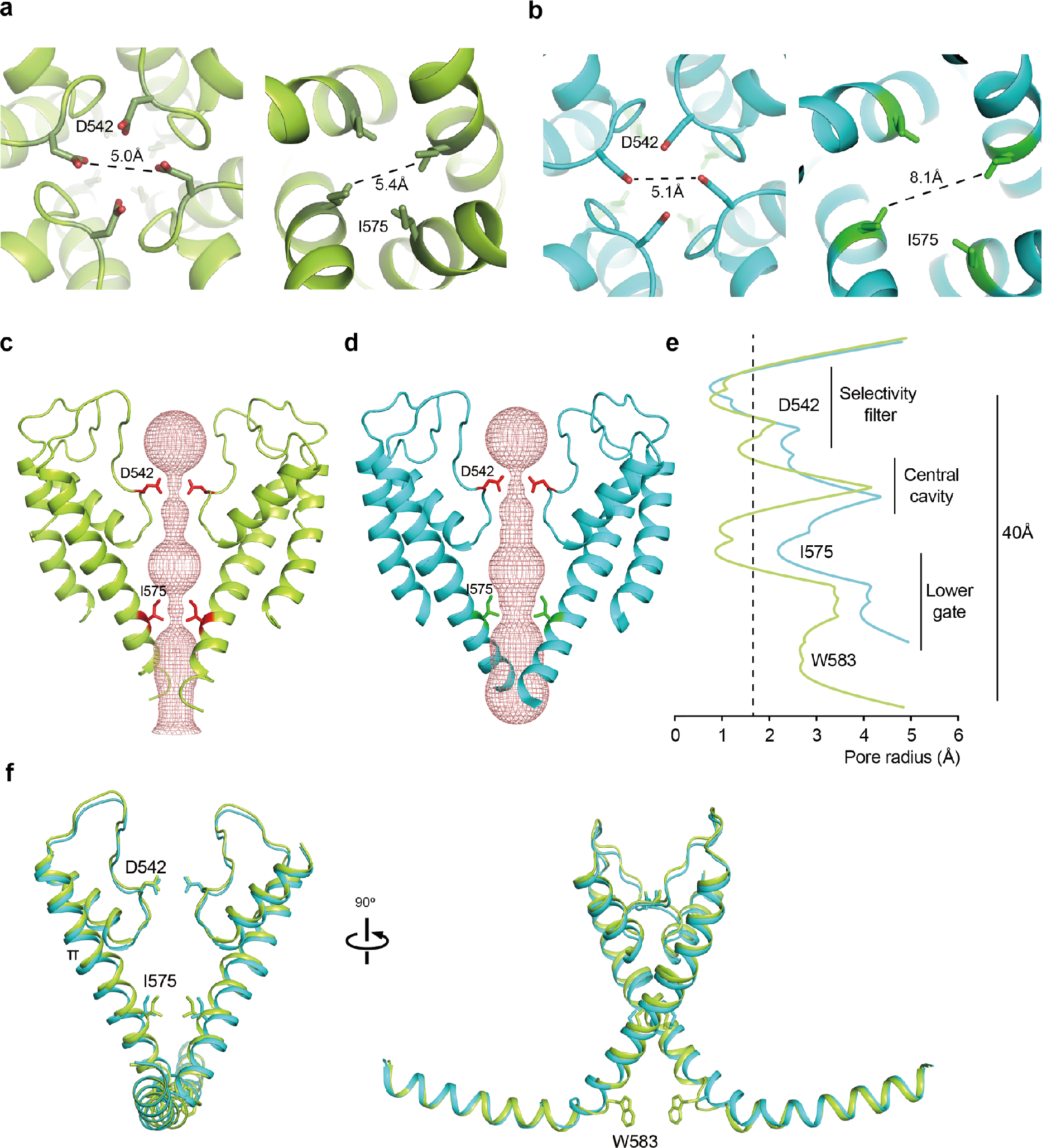
Pore domain characteristics of TRPV5. **a,b,**Top (left) and bottom (right) view of (**a**) TRPV5 FL and (**b**) TRPV5 W583A with subunits individually colored. Side chains are visualized for two constricting residues in the selectivity filter and lower part of the pore, D542 and I575. **c,d,** The ion permeation pathway of (**c**) closed TRPV5 (green) and (**d**) open TRPV5 W583A (blue) is shown as a ribbon diagram with the solvent-accessible space depicted as pink mesh. Only two subunits are shown for clarity, with the side chains of restricting residues (D542 and I575). **e,** Diagram of the pore radius calculated with HOLE is shown for TRPV5 (green) and TRPV5 W583A (blue). The dotted line indicates the radius of a hydrated calcium ion. **f,** Comparison of the TRPV5 (lemon) and TRPV5 W583A (cyan) pore domains with the TRP helix attached. Key residues in the pore domain are depicted.

In contrast to what is seen in the crystal structure of TRPV6(17), there are no significant densities in our structure coordinated by the side chains of T539 or M570, located in the lower part of the pore. Instead we see a weaker density positioned between the four G579 residues, and a second density between the W583 side chains (Supplemental Fig. 7c,e). Both densities are found in the same locations in both half maps and are present when the structure was determined without symmetry, suggesting that they are indeed coordinated cations. W583 is the last residue in the S6 helix, located just before the TRP helix. It is conserved in TRPV6 but not in the other TRPV channels and is essential for TRPV5 function (18) (Supplemental Fig. 10). To explore the influence of W583, we determined a structure of full length TRPV5 with a W583A point mutation in nanodisc to 2.85 Å resolution. Consistent with previously reported ‘gain-of-function’ phenotype for the W583 mutants (18), the lower gate of the TRPV5 W583A structure has an open conformation (Supplemental Fig. 3).

When comparing the pore profiles in both TRPV5 structures, the side chains of D542 form the tightest constriction, just like the corresponding residue D541 in TRPV6. They project into the permeation pathway and partially occlude the pore, creating a minimum interatomic distance between the side chains of 5.0 Å, compared to 5.1 Å in TRPV5 W583A (Fig 3a-d). Analysis of solvent-accessiblity with HOLE revealed a pore radius of approximately 0.7 Å at this position (Fig 3e). Further down, the pore is also tightly constricted at I575, illustrated by a solvent-accessible pore-radius of 1.0 Å (Fig 3e). The interatomic distance between these lower gate side chains is 5.4 Å, approximating the interatomic distances found for the lower gate residues of other TRPV channel structures (Fig. 3a). In contrast, the lower gate of TRPV5 W583A is clearly open for conductance of hydrated calcium ions (Fig. 3b,d,e).

The S6 helix in both wild type and mutant TRPV5 adopts a π-helical conformation (Fig. 3f), which typically results from a single amino-acid insertion that produces a register shift in the protein backbone. From other recent structural studies of TRP channels, such π-helical conformation is suggested to be involved in channel gating (12, 19, 20). Of note, it does not appear to be dependent on sequence, since the residue that induces the register-shift is different in TRPV1 versus TRPV5, but similar between TRPV5 and TRPV6 (Supplemental Fig. 10).

### TRPV5-CaM complex structure

Previous studies suggested that TRPV5 is inactivated in a calcium-dependent manner through the action of CaM (6, 7). CaM consists of N- and C-lobes, each of which contains two calcium-binding sites. Generally, CaM adopts a closed conformation in the calcium-free state and an open conformation in the calcium-bound state. Both states are able to bind various target proteins, including TRP channels, and consequently transduce relevant signals upon calcium binding (21). In the case of modulating TRPV5, the calcium influx induced by channel opening presumably generates calcium-bound CaM that binds to and somehow inactivates the channel.

To gain further insight into this regulatory mechanism, we added bovine CaM to purified and detergent (LMNG) solubilized TRPV5 FL in the presence of 5 mM CaCl, and determined a structure of the complex at 3.0 Å resolution (Fig. 4a,b, Supplemental Fig. 4). We observed major densities at the cytosolic face of the channel that are contributed by the bound CaM (Supplemental Fig. 8). One is located directly under the lower gate and in the symmetry axis; another is located next to the beta sheet and the ankyrin repeats of the two neighboring subunits. To determine the stoichiometry of CaM binding to TRPV5, we performed 3D classification on a density map that was reconstructed with C1 symmetry, and found three different classes. In one class, which contains 35% of particles, one CaM binds to TRPV5 such that its C-lobe interacts with the channel directly below the lower gate, and the N-lobe binds to the beta sheet from inside of the tetramer. In two other classes, however, there are densities corresponding to additional CaMs. In these two classes, one CaM binds to the channel exactly as described above. In addition, there is another N-lobe density that binds to either the opposite or neighboring subunit (Supplemental Fig. 8), indicating that there is a second CaM bound to the channel. The area under the lower gate is surrounded by four TRPV5 subunits and has limited space that allows for binding of only one C-lobe of CaM. Hence, in these latter cases of two CaM bound to one channel, only one CaM binds the channel with both lobes; the second CaM binds only by its N-lobe. Thus, the stoichiometry of CaM binding to TRPV5 is either 1:1 or 2:1. Interestingly, in our reconstruction of full length TRPV5 in nanodisc, we also find extra densities that are similar, but relatively weaker, to the CaM density found in class 1, with one CaM bound to the channel. Most likely, some channels have already bound endogenous CaM in HEK293 cells.

**Figure 4.**
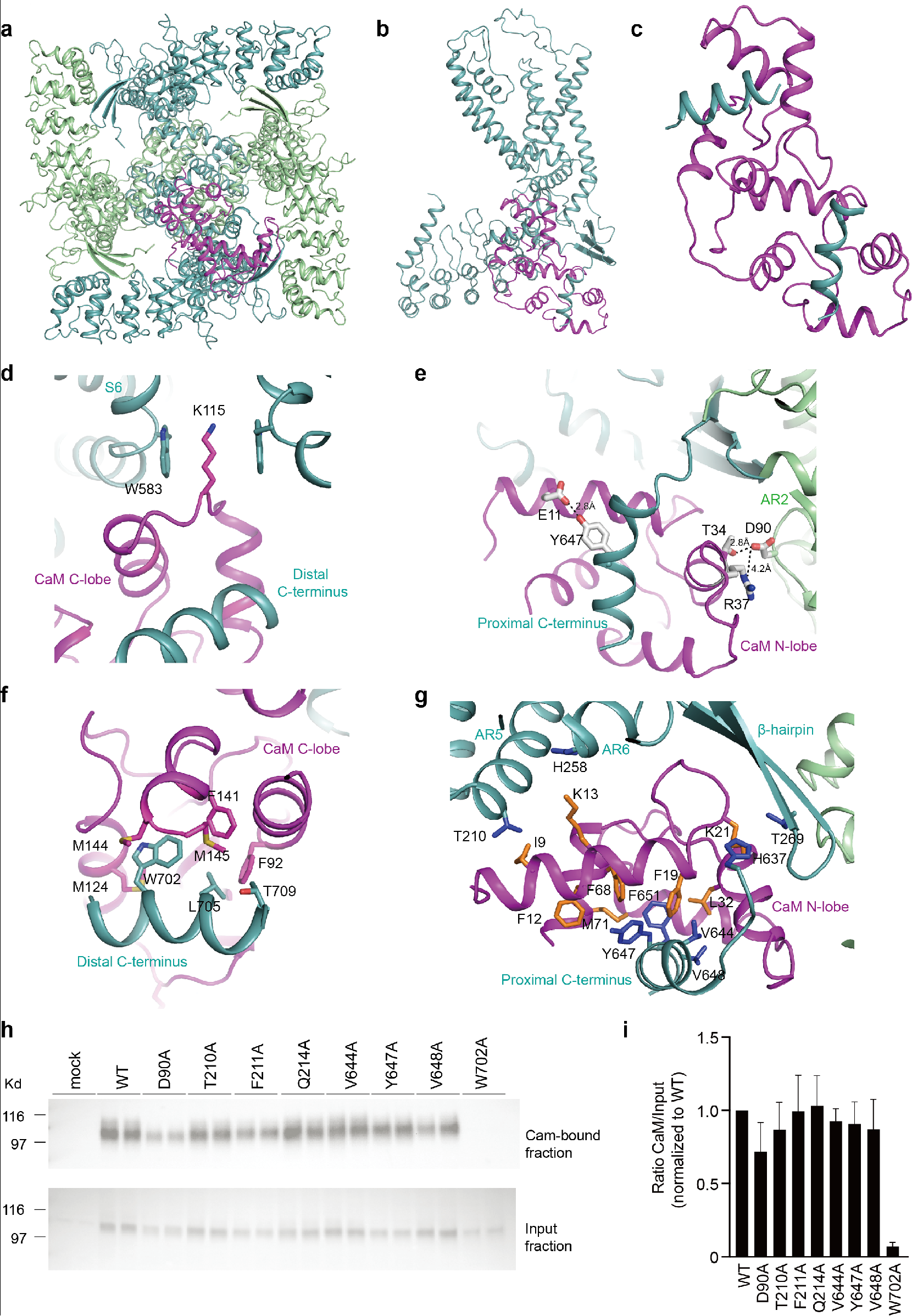
Structure of the TRPV5-CaM complex. **a,** Bottom view of TRPV5-CaM with the two subunits including the CaM-interacting one colored light teal, and the other two in pale green. CaM is colored magenta. **b,** Side view of a TRPV5 monomer (light teal) with CaM (magenta) interaction at the C-terminus. **c,** Overview of CaM (magenta) interacting with two C-terminal helices of TRPV5 (light teal). **d-h** Close-up views of the CaM N-lobe (**e,g**) and CaM C-lobe (**d,f**) interacting with the TRPV5 N- and C-terminus. Side chains of hydrophobic interactions are shown as sticks for both CaM (orange) and TRPV5 (blue). In (**e**) interatomic distances between side chains are depicted. **h,**CaM binding assay of HEK293 cells transfected with either wild type TRPV5 (WT) and the indicated mutants. Samples were analyzed by immunoblotting with GFP antibody. The CaM fraction represents the TRPV5 bound to the CaM agarose beads (top panel) and input demonstrates TRPV5 expression in total cell lysates (bottom panel). Representative immunoblot of three independent experiments is depicted. **i,** Quantification of the immunoblots is depicted as percentage of WT, which represents the relative CaM binding compared to input.

Analysis of the binding interface of TRPV5 and CaM showed that both the CaM N-lobe and C-lobe can bind tightly to TRPV5 at different regions (Fig 4d-g). The CaM N-lobe interacts sandwiched between the beta-sheet and a C-terminal helix (Fig 4e,g). This proximal C-terminal helix (residues 639-653) is not resolved in the apo structures, likely because it is flexible. The binding of CaM N-lobe to the C-terminal helix (639-653) is in line with a previous biochemical finding that the same region in TRPV6, which is highly homologous to TRPV5, can pull down CaM(22). In the TRPV5-CaM complex, this helix is stabilized by CaM and plays an important role in the intermolecular interaction (Fig. 4e,g). Residue Y647 in this helix can form a hydrogen bond with E11 in CaM (Fig. 4e). In addition, four hydrophobic residues in the TRPV5 helix, including V644, Y647, V648 and F651, are surrounded by hydrophobic residues from CaM, which includes F12, F19, K21, L32, F68 and M71 (Fig. 4g). This “V” shaped hydrophobic groove not only stabilizes the helix of TRPV5, but also increases the interaction between TRPV5 and the N-lobe of CaM (Fig. 4g). There is another interaction interface mediated by hydrogen bonds between D90 from the neighboring subunit and T34 in CaM. This interaction was further strengthened by a salt bridge between D90 and R37 in CaM (Fig. 4e). Indeed, our CaM agarose pull-down assay showed that D90A mutation slightly reduces the binding between CaM and TRPV5 (Fig 4h,i). Besides the interface described above, there are three hydrophobic interaction pairs found between TRPV5 and CaM. Residues T210 and H258 of the TRPV5 ARD could form hydrophobic interactions with I9 and K13 in CaM, respectively, and T269 in the β-hairpin can hydrophobically interact with K21 in CaM (Fig. 4g).

The distal C-terminal helix of TRPV5, comprising of residues 698-710, is responsible for CaM C-lobe binding in our structure (Fig 4d,f). The C-lobe of CaM occupies the lumen close to the lower gate such that steric hindrance will preclude binding of a second CaM-lobe. Similar to TRPV6, our TRPV5-CaM complex structure shows that K115 of CaM can insert deeply into the pore region and is surrounded by W583 from four subunits (Fig. 4d) (23). In addition to this interaction mediated by K115 and W583, there is another interaction between the CaM C-lobe and the TRPV5 698-710 helix. The three hydrophobic residues (W702, L705 and T709) in this helix are surrounded by several hydrophobic residues from the CaM C-lobe, including F92, M124, F141, M144 and M145 (Fig. 4f). Of note, the CaM agarose pull-down confirms the previously described interaction of W702 with CaM (Fig 4h,i) (7).

## Discussion

With the current structures, we show that TRPV5 diverted from the thermosensitive TRPV channel permeation mechanics in several ways. First, the pocket analogous to the valilloid binding pocket in TRPV1 is occupied by a regular annular lipid, which is quite different from that of TRPV1, where a phosphatidylinositol binds to the pocket when there is no vanilloid ligand bound to the protein (12). The residues that make contacts between channel and lipid are different (i.e. TRPV5 R460 corresponding to TRPV1 R557). Given that TRPV5 and TRPV6 are considered more stable than their thermosensitive counterparts, the lipid may just provide structural support for normal channel function. Recent data on TRPV6 suggests that R470-lipid interaction is essential for channel conformation as the TRPV6-R470E mutant was solved in a closed state (12).

In addition to the vanilloid pocket, we observed an extensive S1-S2 linker in TRPV5. The position of the linker covers a large portion of the S5-P and P-S6 linkers, thereby shielding them from binding by factors (such as Double Knots toxin in case of TRPV1) from the extracellular space. Besides simply providing a protective blanket against extracellular factor-binding, the S1-S2 linker also connects to the S5-P linker via an intricate interaction network. Preliminary data suggests that this linker is essential for channel function as removal of (part of) the linker results in a non-functional channel (data not shown).

It was suggested that the S6 of TRPV channels interchanges between α-helical and π-helical conformations (12, 19, 20). However, all our TRPV5 structures (closed and open lower gates) were solved with the π-helical conformation. One explanation could be the discrepancy in pore profiles. Despite almost complete sequence conservation with S6, the TRPV structures have different lower gate residues (M577 vs I575). Intriguingly, both TRPV5 and TRPV6 channel have a different lower gate residue, TRPV2 and TRPV6, were solved with their S6 helix in a typical α-helical conformation. On the other hand, the recently published open TRPV6 structure had a π-helical conformation in combination with I575 depicted in the lower gate (12). Future work, carrying out a very focused classification of the S6 region for all available TRPV cryo-EM datasets, may reveal the presence of distinct classes that adopt either a π-helix or α-helix.

Third, TRPV5 and TRPV6 are unique because of their high selectivity for calcium over monovalent ions, which is magnitudes larger than that of the other TRPV channels(1). Because of this feature, TRPV5 and TRPV6 facilitate significant calcium influx at physiological membrane potentials. We showed here that, in contrast to TRPV1 or TRPV2, the conformation of the upper gate in TRPV5 remains unchanged in states where the lower gate is either open and closed. This is in line with the open TRPV6 structure (12), and is consistent with its selective ion permeation. Overall, ion conductance through TRPV5 and TRPV6 resembles that of highly selective voltage-gated cation channels. These types of channels have multiple cations in their selectivity filter, which transiently occupy and are ‘knocked-off’ from sequential cation binding sites. While the TRPV6 crystal structure was solved with multiple calcium binding sites in the selectivity filter (17), only a single density in the selectivity filter domain was observed in our TRPV5 structures. Furthermore, a density lodged between the W583 side chains was found, which does not fit the canonical knock-off model, and needs further investigation. It is likely of significant functional importance, since the TRPV5 W583A mutant was resolved with its lower gate open.

An important feature that TRPV5 shares with many TRP family members and voltage-gated ion channels is CaM-dependent regulation (‘calmodulation’). Currently, little is known about the structural mechanism of CaM regulating TRP channel function. Structural information is mainly limited to structures of CaM bound to peptides of cytoplasmic parts of ion channels.

Our full length TRPV5-CaM complex structure provides novel information about the CaM-dependent channel inactivation and proposes a flexible stoichiometry for TRPV5:CaM binding. Supported by recent NMR studies and the TRPV6-CaM structure (23–25), we now postulate that binding of the CaM C-lobe to the distal C-terminal helix of TRPV5 will close the lower pore, while leaving the proximal C-terminal helix (640-652) of TRPV5 available for binding of a second CaM N-lobe. Since binding of the N-lobe alone is suggested to be much weaker, it might be a dynamic mechanism. In line with the fast channel inactivation, we suggest that the CaM C-lobe binding is likely also very flexible since it is not fixed by interactions provided by TRPV5. Together, this would fit models describing lobe-specific regulation in voltage-gated ion channels (26, 27). The CaM lobes can act as independent calcium sensors based on differential calcium binding affinity (28). This increases its regulatory efficiency and is likely relevant for the vital physiological role of TRPV5 in controlling calcium handling. Within the TRPV subfamily, overlapping CaM binding regions have been suggested for several members despite low sequence homology (22, 29–31). Future studies should clarify whether our proposed mechanism can be extrapolated to other TRPV channels.

In conclusion, our TRPV5 structures, resolved at one of the highest resolutions for TRP channels so far (2.85-3.0 Å), highlight an extended S1-S2 linker that shields the TRPV5 selectivity filter from binding by extracellular factors and provides a rationale why TRPV5 is not activated by vanilloid compounds. Furthermore, the π-helical conformation of S6, visible in both open as well as closed channel states, challenges the idea that all TRPV channels are opened by an α- to π-helical transition of S6. Lastly, our CaM-bound TRPV5 structure increases our understanding of an important calcium-dependent inhibitory mechanism that regulates not only TRPV5 and TRPV6 activity, but also many other TRP channels.

## Methods

### Cloning, expression and purification

Using standard PCR cloning methods, rabbit TRPV5 (rbTRPV5) full length (FL) and residues 1-660 were cloned into a modified BacMam pFastBac1 vector containing an amino-terminal MBP tag separated by a Tobacco Etch Virus (TEV) protease cleavage site. Recombinant bacmid DNA and baculovirus of rbTRPV5 were produced as previously described (32). The plasmid for the membrane scaffold protein (MSP)2N2 was obtained from Addgene (#29520) and was expressed in *Escherichia coli*. It was purified using the N-terminally tagged hexahistidine (His) as described (33). Afterwards, the tag was cleaved with TEV protease and MSP2N2 was snapfrozen and stored at -80 °C.

TRPV5 was purified and reconstituted into nanodiscs in similar fashion to what has been previously described for TRPV1 (9, 34), with minor alterations to the protocol. TRPV5 expression was carried out in HEK293s GnTi- and HEK293F cells, grown in Freestyle 293 Expression medium (Invitrogen) supplemented with 2% fetal bovine serum (FBS) at 37 °C and 8% CO_2_ on an orbital shaker. Cells were transduced with P2 virus when cell density reached 2-2.5 × 10^6^ cells/mL, and sodium butyrate was added 24h after transduction to a final concentration of 10 mM. Cells were collected by centrifugation 48h after transduction, and resuspended in hypotonic buffer (50 mM Tris, 36.5 mM sucrose, 2 mM TCEP, pH 8.0) with freshly added protease inhibitor cocktail (EDTA-Free SigmaFAST tablet, Sigma-Aldrich). Resuspended cells were broken by passing through an emulsifier (Avestin Emulsiflex) five times, and the lysate was cleared by low-speed centrifugation at 8,000x*g* for 20 min at 4 °C. Subsequently, the membrane fraction was collected by ultracentrifugation at 200,000*xg* for 1 hour, and resuspended in buffer A (150 mM NaCl, 2 mM TCEP, 10% glycerol, 50 mM HEPES, pH 8.0). Membranes were solubilized in buffer A supplemented with 20 mM n-dodecyl-β-d-maltoside (DDM, Anatrace) under mild agitation for 2 hours at 4 °C. Detergent-insoluble material was removed by centrifugation at 30,000*xg* for 30 min, and the supernatant was mixed with amylose resin for 1-2 hour to bind the MBP-tagged TRPV5. The resin was washed in a gravity-flow column with 5 column volumes of buffer A supplemented with 10 μg/ml polar soybean lipid mixture and 0.5 mM DDM. The same buffer, with 20 mM maltose added, was used for elution of MBP-TRPV5. The eluted fraction was concentrated to ~1 mg/ml using a 100 kDa Amicon Ultra-4 filter unit.

### Sample preparation

Soybean polar lipids were prepared from a chloroform-dissolved stock solution (Avanti lipids). In short, chloroform was removed from the lipid solution by nitrogen gas drying and subsequent vacuum desiccation for 12 hours. Lipids were rehydrated in buffer A (supplemented with 14 mM DDM) and sonicated, yielding a clear lipid stock at 10 mM concentration.

Reconstitution was performed at two different ratios (TRPV5 monomer:MSP2N2:lipids): 1:1,5:150 for rbTRPV5 660X and 1:2,5:100 for rbTRPV5 FL and rbTRPV5 W583A. The concentrated TRPV5 sample was mixed with soybean polar lipids for 1 hour at 4 °C. Next, MSP2N2 was added and incubated on ice for 5 min, followed by addition of 100 μl of Bio-beads SM2 (Bio-Rad) to remove detergents and initiate nanodisc reconstitution. After rotation for 1 hour at 4 °C, a second batch of Bio-beads (same volume), together with 150 μg of TEV protease was added, and the mixture was rotated overnight at 4 °C. The mixture was filtered to remove all Bio-beads and injected onto a Superdex 200 increase column (GE Healthcare Life Sciences 10/300 GL) in Buffer B (150 mM NaCl, 2 mM TCEP, 50 mM HEPES, pH 8.0).

For the TRPV5-CaM complex, the detergent-solubilized TRPV5 was concentrated to ~1 mg/ml and incubated with bovine calmodulin (Sigma, 1:2 molar ratio of TRPV5 monomer:CaM) and 150 μg TEV protease at 4 °C overnight. The sample was separated on a Superdex 200 increase column (GE Healthcare Life Sciences 10/300 GL) in Buffer B supplemented with 0,02% lauryl maltose neopentyl glycol (LMNG). Of note, 5 mM calcium was added to all buffers.

In all instances, the fractions corresponding to tetrameric TRPV5 were collected and pooled after assessment by SDS-PAGE and negative stain EM. Subsequently, they were concentrated and used for single particle cryo-EM.

### Radioactive calcium uptake assay

To assess the activity of rbTRPV5 FL and 660X, a radioactive ^45^Ca isotope uptake assay was performed as described previously (35). Briefly, HEK293 cells were seeded onto 6-well plates at approximately 1.4×10^6^ cells/well and transiently transfected 6-8h later, with the pcDNA GFP FRT/TO constructs using PEI (polyethylenimine). The next day, cells were trypsinized and re-seeded onto poly-L-lysine-coated 24 wells plates (300 μg poly-L-lysine and 3.0*10^5^ cells per well). Prior to the start of the experiment, cells were pre-treated with 25 μM BAPTA-AM (1,2-bis(o-aminophenoxy)ethane-N,N,N′,N′-tetraacetic acid acetoxymethyl ester) for 30 min. The cells were then washed once with warm KHB buffer (110 mM NaCl, 5 mM KCl, 1.2 mM MgCl_2_, 0.1 mM CaCl_2_, 10 mM Na-acetate, 2 mM NaHPO_4_, and 20 mM HEPES, pH 7.4, with added inhibitors of voltage-gated calcium channels, 10 μM felodipine and 10 μM verapamil), and incubated with KHB buffer supplemented with 1 μCi/ml ^45^Ca for 10 min at 37 °C. Subsequently, the assay was stopped with three washes of ice-cold stop buffer (110 mM NaCl, 5 mM KCl, 1.2 mM MgCl_2_, 10 mM Na-acetate, 0.5 mM CaCl_2_, 1.5 mM LaCl_2_, and 20 mM HEPES, pH 7.4). ^45^Ca uptake was then quantified by liquid scintillation counting.

### Electron-microscopy data acquisition

For negative staining, 3 μl of the purified TRPV5 sample (~0.01-0.1 mg/ml) was applied to glow-discharged EM grids covered by a thin layer of continuous carbon film (TED Pella inc.) and stained with 0.75% (w/v) uranyl formate solution as described (36). EM grids were imaged on a Fei Tecnai T12 microscope (Thermo Fisher Scientific) operated at 120 kV with a TVIPS TemCaM F816 (8K *x* 8K) scintillator-based CMOS camera (TVIPS). Images were recorded at a magnification of 52,000*x*, which resulted in a 2.21 Å pixel size on the specimen. Defocus was set to −1.5 μm.

For cryo-EM, 3 μl of purified TRPV5 sample (~1.0 mg/ml) was applied to glow-discharged holy carbon grids (Quantifoil 400 mesh Cu R1.2/1.3). The grids were blotted by Whatman No. 1 filter paper and plunge-frozen in liquid ethane using a Mark III Vitrobot (FEI) with blotting times of 5-8 seconds at room temperature and 100% humidity. Cryo-EM data collection of TRPV5 took place on a Titan Krios electron microscope (FEI) equipped with a high-brightness field-emission gun (X-FEG) operated at 300 kV at the Howard Hughes Medical Institute Cryo-EM facility at Janelia Research Campus (rbTRPV5 660X), and at UCSF cryo-EM facility (rbTRPV5 FL). Images were collected with a K2-summit direct electron detector (Gatan) using SerialEM in super-resolution mode at a calibrated magnification of x22,500 (0.84 Å physical pixel size) or x29,000 (1.02 Å physical pixel size) for TRPV5 660X and TRPV5 FL respectively. Dose rate, total dose and defocus range used for data collection were summarized in supplementary table 1.

### Data processing

For negative-stain data, RELION was used for particle picking and 2D classification.

For cryo-EM data, the 60-frame image stacks were drift-corrected and 2×2 binned by Fourier cropping using MotionCor2 (37). All micrographs were converted to PNG-image format and individually inspected by eye. Motion-corrected sums without dose-weighting were used for contrast transfer function estimation with GCTF (38). Motion-corrected sums with dose-weighting were used for all other image processing.

Dataset of TRPV5 660X, TRPV5 FL in LMNG or lipid nanodiscs were processed with similar strategy. In summary, automatically picked particles by Gautomatch (http://www.mrc-lmb.cam.ac.uk/kzhang/Gautomatch/) were extracted by RELION and then imported into cryoSPARC (39) for 3D classification and refinement procedures. The initial model was generated using *ab-initio* reconstruction. Particles from good classes by *ab-initio* reconstruction were further sorted by iterative 2D classification and heterogenous refinement. Well-sorted particles were finally subjected to homogenous refinement to generate the final maps, including two half maps, unsharpened map and sharpened map. Resolution was estimated using FSC=0.143 criterion. Local resolution estimates were calculated with the unsharpened raw density map using ResMap. Detailed information of data processing for the TRPV5 660X and TRPV5 W583A datasets can be found in Supplementary figure 5.

### Model building

*Ab-initio* model building was carried out in COOT and PHENIX (40, 41). The initial model was generated with SWISS-MODEL on the basis of sequence alignment of TRPV5 with crystal structure of TRPV6 (PDB:5IWK). The resulting model was refined in real space with Phenix.real_space_refine and subsequently adjusted manually in COOT. This process was repeated until satisfaction of the Ramachandran validation. Further validation was carried out using EMRinger and MolProbity in PHENIX. The program HOLE was used to calculate the pore profile of TRPV5 (42). Chimera (43) and Pymol were used to prepare images.

## Data availability

Cryo-EM density maps of TRPV5 have been deposited in the Electron Microscopy Data Bank (EMDB) under the accession numbers EMD-XXXX, EMD-XXXX, EMD-XXXX and EMD-XXXX. Particle image stacks after motion correction related to TRPV5 have been deposited in the Electron Microscopy Public Image Archive (http://www.ebi.ac.uk/pdbe/emdb/empiar/) under accession numbers EMPIAR-XXXXX (for C-terminal truncated TRPV5 660X reconstituted in nanodiscs), EMPIAR-XXXXX (for full length TRPV5 reconstituted in nanodiscs), EMPIAR-XXXXX (for complex of TRPV5 and CaM in LMNG) and EMPIAR-XXXXX (for TRPV5 with W583A mutant reconstituted in nanodiscs). Atomic coordinates for TRPV5 have been deposited in the PDB under the accession numbers XXXX, XXXX, XXXX and XXXX. All electrophysiological data generated or analyzed during this study are included in the published article. All other data are available from the corresponding authors upon reasonable request.

## Acknowledgements

The authors thank N. Thijssen for technical support, and C. Hong, R. Huang and Z. Yu at the HHMI Janelia Cryo-EM Facility for help with data acquisition. We also thank D. Bulkley, A. Myasnikov and M. Braunfeld at UCSF cryo-EM facility for help with data acquisition. J.W. is supported by the European Union’s Horizon 2020 Marie Skłodowska-Curie (grant agreement No 748058) and by The Netherlands Organisation for Health Research and Development (Off Road grant 451001 004). S.D. is supported by a Human Frontier Science Program (HFSP) Postdoctoral Fellowship. M.G. was supported by a Dutch Kidney Foundation student abroad fellowship and by the Nora Baart Foundation (NBS) of the Netherlands Foundation for the Advancement of Biochemistry (NVBMB). Y.C. is Investigator with the Howard Hughes Medical Institute.

## Contributions

All authors designed experiments. S.D., M.G., and J.W. expressed and purified all protein samples used in this work and J.W. performed the functional analysis. S.D. carried out all cryo-EM experiments, including data acquisition and processing. S.D. build the atomic model based on the cryo-EM maps. D.J., Y.C. and J.W. supervised experiments and data analysis. All authors analyzed data and wrote the manuscript.

